# Host cell maturation modulates parasite invasion and sexual differentiation in *Plasmodium*

**DOI:** 10.1101/2021.07.28.453984

**Authors:** Franziska Hentzschel, Matthew P Gibbins, Charalampos Attipa, Dario Beraldi, Christopher A Moxon, Thomas D Otto, Matthias Marti

## Abstract

Malaria remains a global health problem with over 400,000 deaths annually^1^. *Plasmodium* parasites, the causative agents of malaria, replicate asexually in red blood cells (RBCs) of their vertebrate host, while a subset differentiates into sexual stages (gametocytes) for mosquito transmission. Parasite replication and gametocyte maturation in the erythropoietic niches of the bone marrow and spleen contribute to pathogenesis and drive transmission^2^, but the mechanisms underlying this organ enrichment remain unknown. We performed a comprehensive single cell analysis of rodent *P. berghei* in spleen, bone marrow and blood to define parasite phenotypes specific to those niches. Single cell RNA-seq analysis of host and parasite cells reveals an interferon-driven host response to infection as well as transcriptional adaptations of *Plasmodium* to RBC maturation status. We show that *P. berghei* exhibits a bimodal invasion pattern into either normocytes or early reticulocytes and, using functional assays, identify CD71 as a host receptor for reticulocyte invasion. Importantly, we observe an increased rate of gametocyte formation in reticulocytes that is nutrient-dependent and triggered post invasion (i.e., same cycle sexual commitment). Our data provides a thorough characterisation of host-parasite interactions in erythropoietic niches and defines host cell maturation state as the key driver of parasite adaptation.

## Main

The erythropoietic niches in the extravascular parenchyma of bone marrow and spleen are unique sites of *Plasmodium* asexual and sexual development^2–4^. During erythropoiesis, erythroblast precursors enucleate to give rise to reticulocytes, which intravasate into circulation and further mature to normocyte RBCs. This process is accompanied by sequential shedding of surface proteins such as CD44 and the transferrin receptor CD71, and loss of RNA (Extended Fig. 1a)^5,6^. Recent studies indicate specific parasite adaptations to host organ and host RBC maturation status, including cryptic parasite replication cycles and increased gametocyte formation^3,7–13^. Yet, deciphering the mechanistic basis for those phenotypes has been hampered by the complex cellular composition of erythropoietic niches, which is difficult to disentangle in bulk approaches like RNA sequencing (RNA-seq). In contrast, advances in single cell profiling have enabled the analysis of thousands of individual non-synchronous cells in parallel^14,15^. Using scRNA-seq, flow cytometry and functional assays, we set out to define *Plasmodium* and host-specific phenotypes in blood and erythropoietic niches at single cell level and to elucidate the contribution of both host organs and cells to parasite behaviour.

### Quantitative analysis of *P. berghei* infection in blood and erythropoietic tissues

To obtain an initial quantitative understanding of parasite distribution across host cells and organs, we followed the course of a *P. berghei* blood stage infection over three days by flow cytometry. Gating on CD45^-^/Ter119^+^ RBCs in spleen, bone marrow and blood, we used CD71 as marker for reticulocytes and CD44 to differentiate extravascular CD71^+^/CD44^+^ reticulocytes from intravascular CD71^+^/CD44^-^ reticulocytes (Extended Data Fig. 1a-c)^6,7^. At all three time points, parasitaemia was highest in spleen and lowest in bone marrow (Fig. 1a, Extended Data Fig. 1d) and especially splenic and blood intravascular reticulocytes were highly parasitized (Fig. 1a). Note that, although blood CD71^+^/CD44^+^ RBCs are highly infected, their overall number in blood is very low (Fig. 1b, Extended Data Fig. 1b, c). We calculated the relative enrichment of host cell types in iRBCs per organ as the log-fold change of host cell proportion in iRBCs over host cell proportion in all RBCs (Fig. 1b). In spleen and blood, intravascular CD71^+^/CD44^-^ reticulocytes were enriched in iRBCs, supporting the *P. berghei* preference for reticulocytes. In contrast, extravascular CD71^+^/CD44^+^ reticulocytes in spleen and bone marrow were only invaded at later stages of infection, indicating that in initial days of the infection, *P. berghei* does not have access to the extravascular erythropoietic niches. The increased infection rate of extravascular reticulocytes at day 3 (spleen) and 4 (bone marrow) coincides with reported vascular leakage in those organs^10^. In line with these observations, sections of *P. berghei*-infected spleens showed histopathological signs of major splenic damage, including increasing levels of haemorrhage, necrosis, inflammation, as well as white pulp hyperplasia and lesions at later stages of infection (Fig. 1c, Supplementary Table 1). Immunofluorescence staining of spleen sections additionally confirmed initial parasite restriction to the vasculature, with extravascular parasites only detected 3 to 4 days post infection (dpi) as the vasculature becomes more disrupted and fragmented (Fig. 1d, Extended Fig. 2).

**Figure 1:**
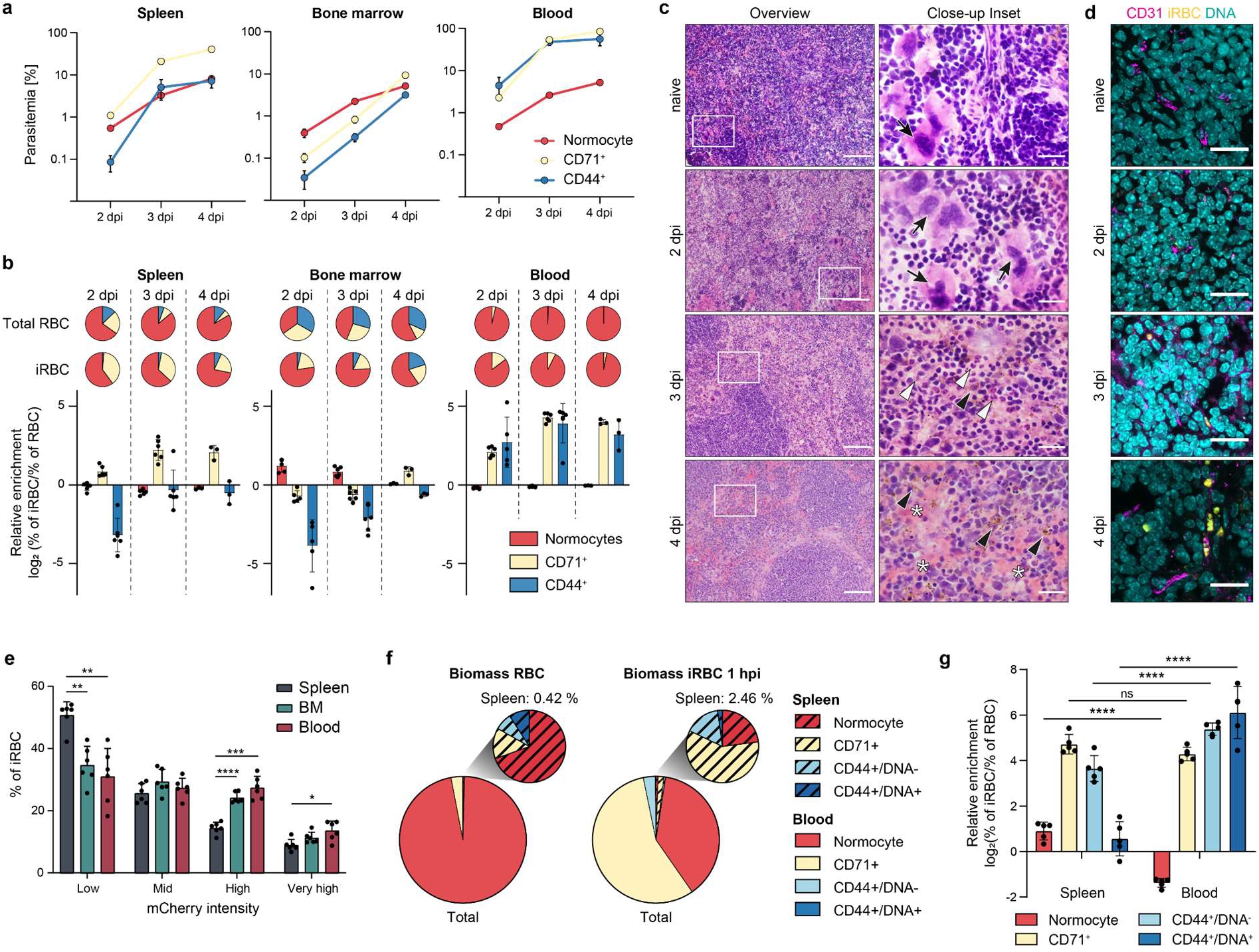
*P. berghei* host cell preference in spleen, bone marrow and blood. **a**, Parasitemia per host organ and cell after infection with 1*10^6^ iRBC i.v. (n = 6 except for 4 dpi, where n = 3). **b**, Pie charts: average cell composition of RBCs (upper row) and iRBCs (lower row) per day and organ. Bar plots: Relative enrichment of host cell types in iRBC versus RBCs per organ. Values > 0 indicate preferential invasion of this host cell type, values < 0 indicate decreased invasion. Same groups as in a. **c**, Representative histology images of naive and infected spleen. Left column: Scale bar = 100 μm. Box indicates area of close-up. Right column: Scale bar = 20 μm. Black arrow: megakaryocytes indicative of hematopoiesis; white arrow: necrosis; black arrowhead: pigmented macrophages; white arrowhead: neutrophils, asterix: haemorrhage. **d**, Representative immunofluorescence images of spleen sections stained for CD31 (magenta). Parasites are mCherry-fluorescent (pseudocolored yellow). Scale bar = 20 μm. **e**, Percentage of parasites per mCherry gate per organ as proxy for parasite stages 3 dpi. (n = 6. 2-way ANOVA, Tukey’s post test). **f**, Biomass of RBC cell types (left) and iRBC cell types 1 hpi with 4*10^7^ iRBC (right) in total RBCs (large pie chart) and spleen RBCs (small pie chart) (n = 5). **g**, Relative enrichment of host cell types in iRBC versus all RBC. Values > 0 indicate preferential invasion of this host cell type, values < 0 indicate decreased invasion. (n = 5. 2-way ANOVA, Sidáks post test). **a**, **b**, **e**, **g**: Mean +/- SEM. *, p < 0.05, **, p < 0.01, ***, p < 0.001, ****, p < 0.0001.

Using the 5’hsp70-driven mCherry reporter as a proxy for parasite maturation^16^, we found young parasite stages enriched in the spleen (Fig 1e, Extended Data Fig. 1b, e). This enrichment could either be the result of preferential RBC invasion in the spleen or of retention of young iRBCs during splenic passage, as observed for *P. falciparum* ring stages^17^. To differentiate between these two scenarios and capture parasite localisation immediately after invasion, mice were infected with synchronised schizonts and ring stage distribution was quantified 1 hour post invasion (hpi). We focussed on the analysis of spleen and blood, as bone marrow showed low infection rates and thus likely does not contribute much to parasite biomass at invasion (Fig. 1a, Extended Fig. 1d). A DNA stain enabled us to differentiate enucleated CD71^+^/CD44^+^/DNA^-^ reticulocytes from nucleated CD71^+^/CD44^+^/DNA^+^ erythroblasts (Extended Data Fig. 1a, 3a).

Parasitaemia was approximately 10-fold higher in CD71^+^/CD44^-^ reticulocytes compared to normocytes, while splenic erythroblasts were infected even less frequently than normocytes (Extended Fig. 3b). Estimation of total biomass revealed that only about 2.4 % of all parasites were found in the spleen (Fig. 1f), much lower than what has been observed for *P. vivax^3,4^*. Despite the apparent tropism for reticulocytes, 37.87 % of all parasites were present in CD71^-^ normocytes in the blood, demonstrating that *P. berghei* is not restricted to invasion of reticulocytes (Fig. 1f, Extended Data Fig. 3f). Blood CD71^+^/CD44^+^/DNA^-^ and DNA^+^ cells were highly parasitized, but due to their low prevalence (below 0.1 % RBCs) they contributed little to the overall parasite biomass (Fig. 1f). While splenic RBCs had an approximately 5-fold higher chance to be infected than circulating RBCs, this preference was mainly due to the high abundance of reticulocytes in this organ, and the parasite tropism for CD71^+^ cells was independent of the organ (Fig. 1g). Interestingly, normocyte infection rates were also higher in spleen compared to blood, potentially because of the local accumulation of schizonts in the splenic sinusoids before rupture (Fig. 1g). These findings are largely independent of starting parasitaemia, as similar results were obtained using a 10-fold lower inoculation dose (Extended Data Fig. 3c-f). In summary, *P. berghei* preferentially invades intravascular reticulocytes in spleen and blood, yet a substantial proportion of parasites is found within normocytes. In contrast, extravascular RBCs only become accessible to the parasite following infection-induced vascular damage.

### Single-cell RNAseq reveals interferon-driven host response to infection

We hypothesised that *Plasmodium* parasites would respond to the different environmental conditions encountered in these diverse host organs and cells and thus investigated transcriptional changes of parasite and host cells in the different compartments. To this end, we enriched for *P. berghei*-infected splenic, bone marrow and blood cells by flow sorting, labelled surface CD71 and CD44 expression with barcoded antibodies (Cellular Indexing of Transcriptomes and Epitopes by Sequencing, CITE-seq^18^) and analysed host and parasite transcriptomes by droplet-based scRNA-seq (Fig. 2a, Extended Data Fig. 4)^19^. After removal of low-quality cells, the dataset consisted of 14,953 spleen, 13,509 bone marrow and 12,973 blood cells from two mice, which included a total of 13,128, 7,132 and 12,960 *P. berghei*-infected cells, respectively (Supplementary Table 2). Additional 2,746 spleen, 8,500 bone marrow and 66 blood cells were obtained from one uninfected control mouse (Supplementary Table 2). Importantly, for all samples, the mean number of RNA transcripts (unique molecular identifier, UMI) and mean number of genes detected per cell were comparable to or higher than in the previously published malaria cell atlas (Supplementary Table 2).

**Figure 2:**
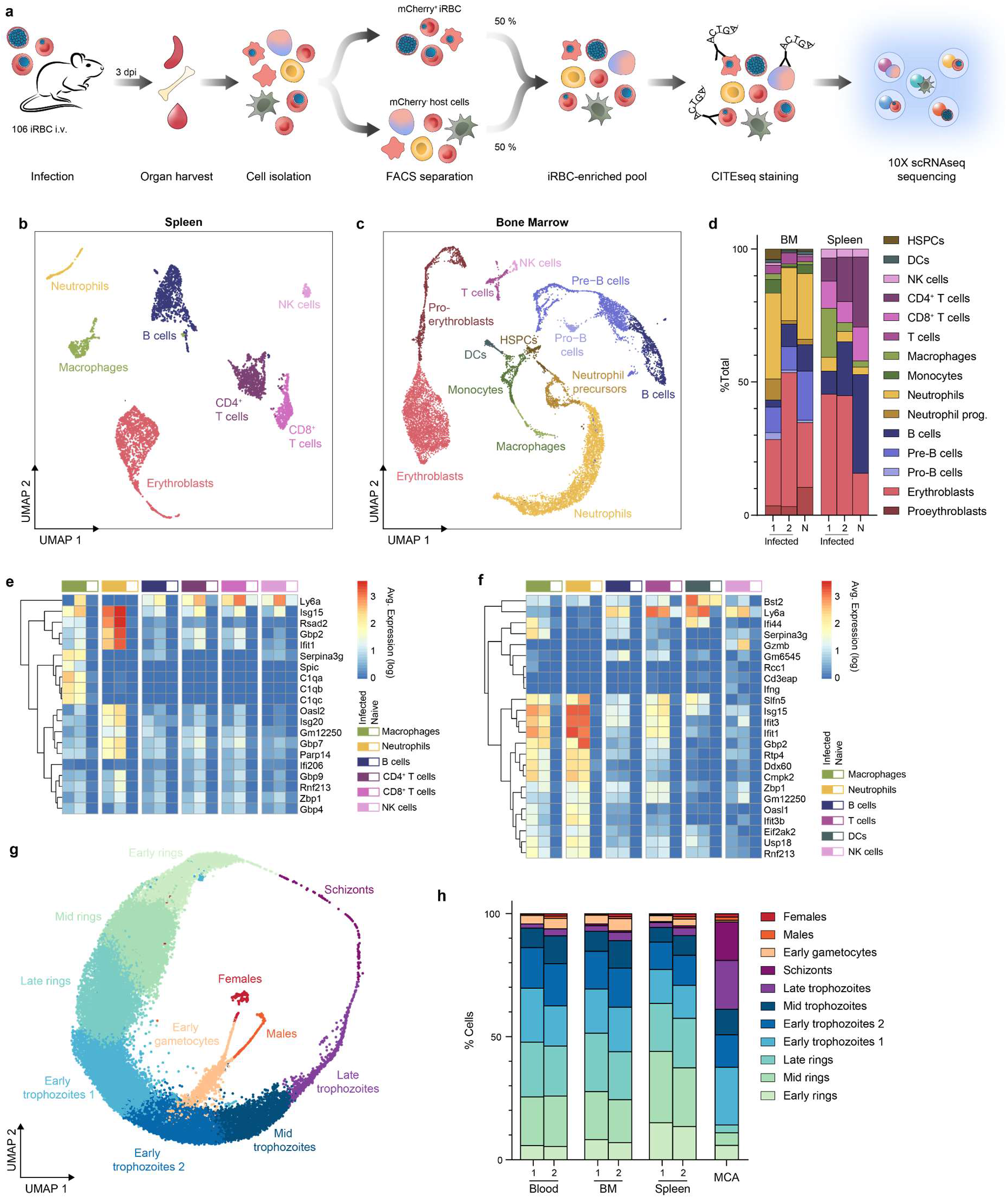
scRNA-seq analysis of *P. berghei* and host cells from spleen, bone marrow and blood. **a**, Cartoon depicting experimental strategy to enrich for infected RBCs derived from spleen, bone marrow and blood for subsequent scRNA-seq analysis. **b**,**c**, UMAP of spleen (**b**) and bone marrow (**c**) host cells, colored according to clusters. **d**, Cell type distribution in infected and naïve (N) spleen and bone marrow host cells (infected: n = 2, naïve: n = 1). **e**, **f**, Heatmap showing average expression level of genes upregulated upon infection across cell types in the spleen (**e**) and bone marrow (**f**). All genes plotted were among the five most significantly upregulated genes in at least one cell type. **g**, UMAP of *P. berghei* cells. **g**, Relative proportion of parasite clusters across organs. MCA: malaria cell atlas, BM: bone marrow. (n = 2 per organ, MCA n = 1)

Graph-based clustering of splenic and bone marrow host cell transcriptomes followed by annotation based on marker gene expression and public reference datasets^14,20^ identified seven (spleen) and thirteen (bone marrow) distinct cell types that were visualised by uniform manifold approximation and projection (UMAP) (Fig. 2b, c, Extended Fig. 5a, b, Supplementary Table 3). Splenic clusters included CD4^+^ and CD8^+^ T cells, mature B cells, macrophages, neutrophils, erythroblasts and NK cells. Bone marrow clusters consisted of immature and mature B cells, HSPCs, T cells, NK cells, proerythroblasts and erythroblasts, dendritic cells, monocytes, macrophages, neutrophil precursors, and neutrophils (Fig. 2b, c). Proportions of cell types did not differ significantly between infected and uninfected mice. However, a trend for an expansion of erythroblasts in infected spleens might indicate stress erythropoiesis (Fig. 2d).

*Plasmodium* infection caused a major dysregulation of gene expression in immune cells in bone marrow and spleen, and the majority of significantly differentially expressed (FDR < 0.01) genes were upregulated upon infection (Extended Data Fig. 6a, b, Supplementary Table 4). Many of those genes were shared across cell types and included genes that are part of the interferon response, such as Isg15, Ifit1 and 3 or Ly6a (Fig. 2e, f). Notably, splenic macrophages also upregulated genes of the complement system, such as C1qa, C1qb and C1qc. Gene ontology (GO) analysis confirmed the induction of a strong type I interferon and interferon γ response in bone marrow and spleen (Supplementary Table 5). Only few genes were downregulated upon infection, with no notable GO term enrichment (Supplementary Table 5). Overall, host response to parasite infection in the spleen and bone marrow was governed by a strong interferon-driven response with minor cell-type specific variations.

### Identification of CD71 as a host cell receptor for *P. berghei* invasion into reticulocytes

To assign developmental stages to the parasite single cell transcriptomes, we integrated the *P. berghei* data sets from spleen, bone marrow and blood with the malaria cell atlas (MCA)^15^, yielding twelve clusters (Extended Data Fig. 7a). Correlation with the MCA and identification of conserved marker genes enabled annotation of eleven clusters, while one cluster (cluster 7) with ambiguous marker genes and very low transcripts per cell was excluded as putative artifact (Extended Data Fig. 7b-d, Supplementary Table 6). Using UMAP visualisation, most parasite cells arranged in circular fashion following the asexual cycle, whereas one cluster corresponding to early gametocytes diverged from early trophozoites and lead to female and male gametocyte clusters (Fig. 2g). The dataset is enriched towards ring and early trophozoite stages because of the early time point of the harvest and the semi-synchronicity of *P. berghei* infections (Fig. 2g, h). Confirming the flow data, spleen harboured more early ring stages than bone marrow and blood (Fig. 2h, Extended Data Fig. 7e). Comparing profiles across organs, the only notable differential gene expression was detected in late trophozoites, where late-stage markers such as MSP1, SERA1, RON4 and MSRP2 are downregulated in blood (Extended Data Fig. 7f, Supplementary Table 7).

These differences probably reflect the sequestration of more mature trophozoites and schizonts in spleen and bone marrow compared to blood^21^.

Next, we aimed to classify the infected host cells in order to define host-cell dependent differences in parasite transcriptomes. CD71 surface expression (as determined by CITE-seq) was exclusive to very early rings and was rapidly lost in further developmental stages, suggesting reticulocyte co-maturation with parasite development (Fig. 3a). Reticulocyte maturation to normocytes is also accompanied by a loss of RNA content^5^. Indeed, using the number of detected host transcripts (*Mus musculus* UMI, MmUMI) as a proxy for host cell RNA, we found that CD71^+^ reticulocytes (CD71 signal > 1.5) contained a high amount of RNA (Fig. 3b). In line with previous observations, loss of CD71 surface expression preceded host RNA degradation (Fig. 3b)^5^. We thus used MmUMI as a sensitive marker to determine RBC maturation status, and defined host cells with > 100 MmUMI as reticulocytes and cells with < 100 MmUMI as normocytes (Fig. 3c).

**Figure 3:**
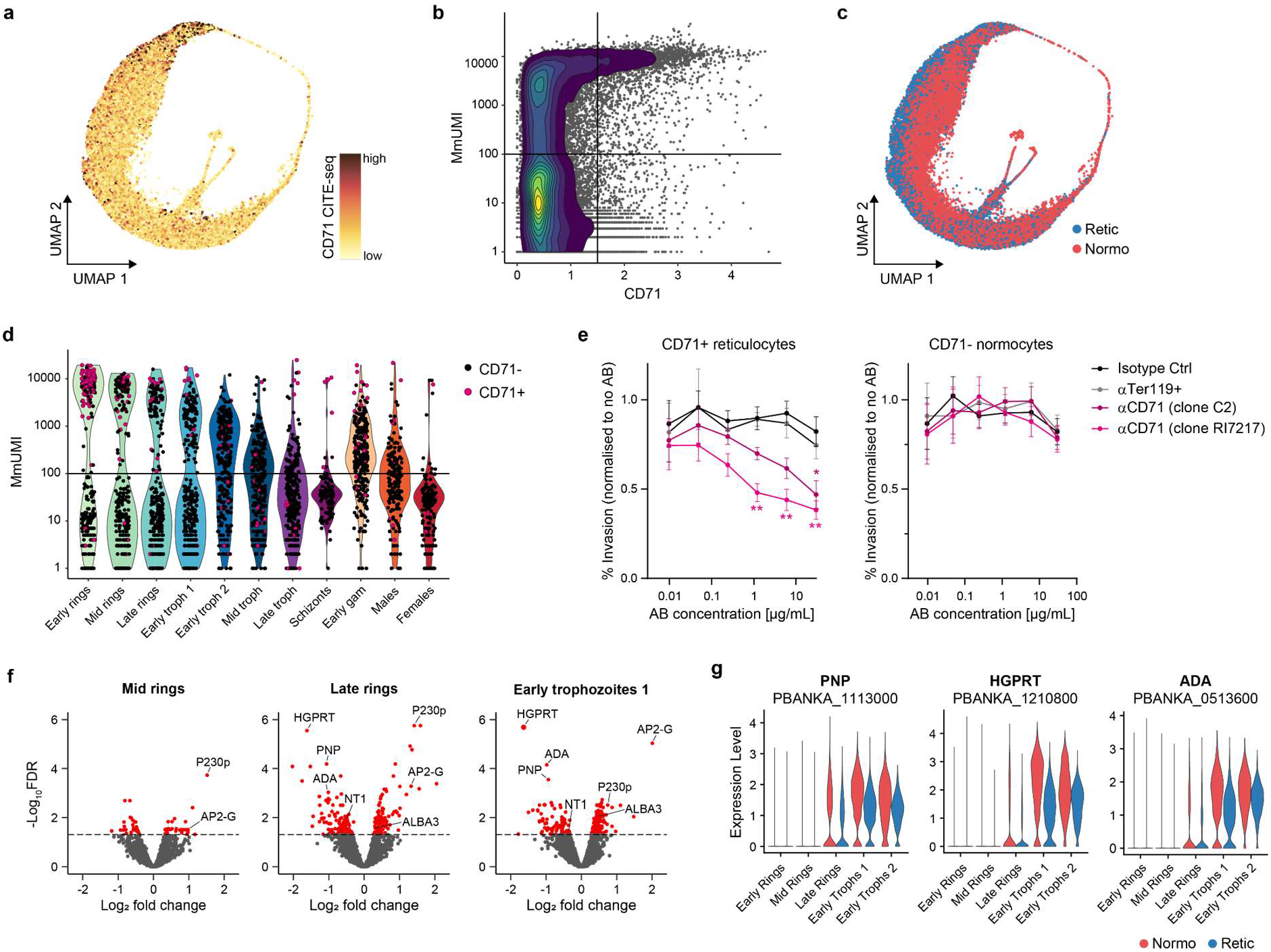
*P. berghei* adapts to its host cell maturation status. **a**, UMAP of *P. berghei* cells colored by CD71 surface protein levels as measured by CITE-seq. **b**, Host RNA levels, quantified as mouse-derived UMI (MmUMI), plotted against CD71 surface protein levels as measured by CITE-seq. Horizontal and vertical lines indicate thresholds separating CD71-positive (CD71 > 1.5) from CD71-negative cells and reticulocytes (MmUMI > 100) from normocytes, respectively. **c**, UMAP of *P. berghei* cells colored by host cell (red: normocytes, blue: reticulocytes). **d**, Host RNA levels across stages. Cells are randomly subsampled to 300 cells per stage. Colour of dots indicates CD71 positive (pink) and negative cells (black). **e**, Invasion of merozoites in CD71-positive reticulocytes (left panel) or CD71-normocytes (right panel) in the presence of anti-CD71 antibody (2 different clones), anti-Ter119 antibody or isotype control. Invasion was normalised to no antibody (AB) treatment and significance tested against isotype control. (n = 6. Mean +/- SEM. 2-way ANOVA, Dunnets post test. *, p < 0.05, **, p < 0.01). **f**, Volcano plot of differential gene expression in reticulocyte-infecting parasites versus normocyte-infecting parasites in selected stages. Red: genes passing the false discovery rate (FDR) threshold. **g**, Expression level of genes of the purine salvage pathway in early parasite stages, separated by host cell (red: normocytes, blue: reticulocytes). UMI: Unique molecular identifier, Retic: reticulocytes, Normo: normocytes. AB: Antibody. FDR: false discovery rate.

Host RNA content decreased with parasite development, further supporting co-maturation of parasite and host cell (Fig. 3d). Notably, early rings were only detected in reticulocytes with a high RNA content (> 1000 MmUMI) or in normocytes (Fig. 3d). The lack of early rings in reticulocytes with intermediate RNA content (100 < MmUMI <1000) suggested a bimodal invasion pattern based on availability of specific host cell receptors. As most reticulocytes harbouring early rings were also CD71^+^ (Fig. 3d), and recent work has revealed that CD71 is an entry receptor for the human malaria parasite *P. vivax^22^*, we hypothesised that CD71 might also be a receptor for *P. berghei* reticulocyte invasion. Indeed, two different monoclonal antibodies targeting CD71 blocked *P. berghei* invasion into reticulocytes in a dose-dependent manner but did not inhibit invasion of normocytes (Fig. 3e, Extended Fig. 8), indicating that *P. berghei* requires CD71 as receptor for reticulocyte invasion, yet utilizes an alternative receptor for normocyte invasion.

### Host cell maturation triggers sexual differentiation in *P. berghei*

Intriguingly, parasite cells separated on the UMAP according to their host cell, indicating transcriptional adaptation to the different host environments (Fig. 3c). We analysed differential gene expression in early rings to early trophozoites as in those stages, normocyte and reticulocyte host cells could be clearly differentiated (Fig. 3d). Indeed, signatures of differential gene expression depending on the host cell were detected in mid ring, late ring and early trophozoite 1 clusters (Fig. 3f, Supplementary Table 8). Specifically, purine nucleoside phosphorylase (PNP, PBANKA_1113000), hypoxanthine-guanine phosphoribosyltransferase (HGPRT, PBANKA_1210800), adenosine deaminase (ADA, PBANKA_0513600), and the nucleotide transporter 1 (NT1, PBANKA_1360100) were upregulated in normocyte-infecting parasites (Fig. 3f, g). These genes are all part of the purine salvage pathway to synthesise purine nucleotides from host metabolites and their upregulation might compensate for the limited abundance of those metabolites in normocytes compared to reticulocytes^23^.

One of the most notable genes upregulated in reticulocyte-infecting parasites was AP2-G, a transcriptional master switch activating the first set of sexual stage genes during sexual commitment (Fig. 3f)^23,24^. Also other gametocyte-related genes such as ALBA3 and P230p^24^ exhibited higher expression in reticulocyte-infecting parasites, supporting a link between sexual commitment and reticulocyte invasion (Fig. 3f). On the UMAP visualisation, AP2-G-positive cells were most prevalent in early ring to early gametocyte stages and appeared to follow reticulocyte distribution (Fig. 4a, Fig. 3c). Overall AP2-G expression levels per parasite stage did not differ between normocyte- and reticulocyte-infecting parasites (Fig. 4b). However, the proportion of early parasite stages positive for AP2-G was significantly higher in reticulocytes compared to normocytes, independent of host organ (Fig. 4c), indicating that the host cell, not the organ modulates sexual commitment.

**Figure 4:**
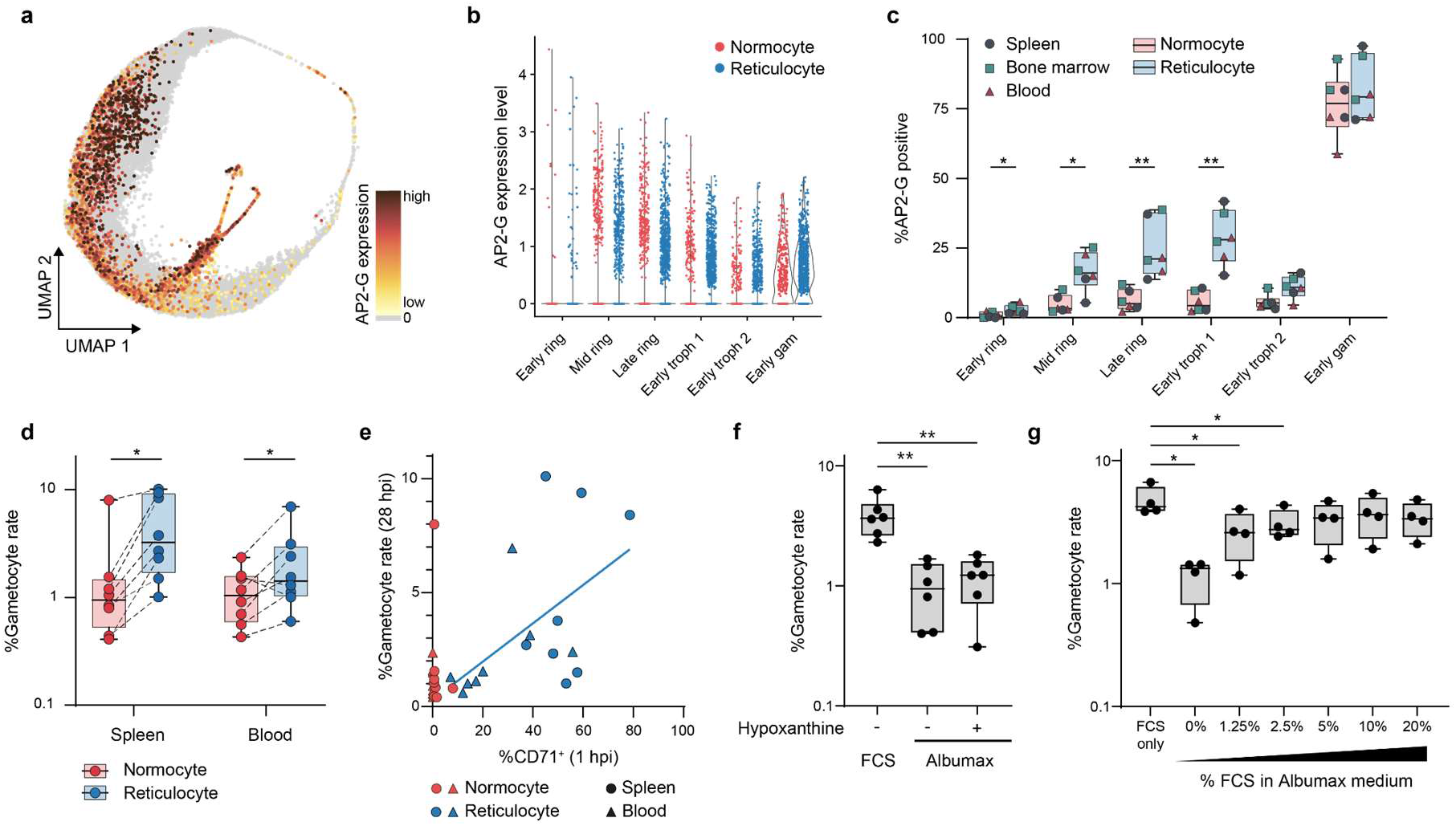
*P. berghei* sexual development occurs preferentially in reticulocytes. **a**, UMAP of *P. berghei* coloured by AP2-G expression strength. **b**, AP2-G expression level in early stages, separated by host cell. **c**, Proportion of AP2-G positive cells in early stages, separated by host cell (box plot colour). Individual dots are coloured by organ. (n = 6). **d**, Gametocyte conversion rate after 28 h *ex vivo* culture of normocyte- or reticulocyte-enriched iRBC isolated from spleen or blood. Samples from one organ were matched across host cell. (n = 8. Two-way ANOVA, Tukey’s post test). **e**, Correlation between proportion of CD71^+^ cells 1 hpi and subsequent gametocyte rate after 28 h *ex vivo* culture. Same samples as in d. **f**, Gametocyte rate after *ex vivo* culture in nutrient-rich or -poor medium. (n = 6. One-way ANOVA, Dunnett’s post test.) **g**, Gametocyte rate after *ex vivo* culture in minimal medium supplemented with varying concentrations of FCS. (n = 4. One-way ANOVA, Dunnett’s post test). **c**, **d**, **f**, **g**, Center line, median; box limits, upper and lower quartiles; whiskers, min to max range; all points shown. *: p value < 0.05; **: p value < 0.01.

To independently investigate this observation, we quantified gametocyte rates in different host cells *ex vivo*. Using a gametocyte reporter line (PbGFP_CON_/RFP_GAM_)^25^, we magnetically sorted freshly invaded RBCs into a CD71^+^-enriched reticulocyte and a CD71-depleted normocyte fraction and quantified the proportion of parasites that developed into gametocytes (gametocyte rate) after 28 h *ex vivo* culture by flow cytometry (Extended Data Fig. 9a-e). As expected, parasitaemia was higher in reticulocytes compared to normocytes, with a specific enrichment of gametocytes (Fig. 4d, Extended Data Fig. 9f). Gametocyte rates in the reticulocyte fraction correlated with the proportion of CD71^+^ cells after purification (Pearson, r = 0.53, n = 16, p = 0.0332), although the latter might be underestimated due to steric hinderance of CD71 by the anti-CD71 antibodies used for magnetic cell purification (Fig. 4e). Notably, there was no significant difference in the gametocyte rate between spleen and blood (Fig. 4d). In summary, both scRNA-seq and *ex vivo* data indicate that sexual commitment is higher in reticulocytes than in normocytes, independent of host organ.

We hypothesised that purine levels serve as metabolic triggers for sexual commitment, since downregulation of the purine salvage pathway has been previously correlated with a decrease in gametocytemia^26^. However, the addition of the purine precursor hypoxanthine to minimal medium containing Albumax II did not affect the basal gametocyte rate of about 1 % (Fig. 4f). In contrast^23^, when culturing young ring stages in nutrient-rich standard medium supplemented with foetal calf serum (FCS), we saw a striking increase of sexual commitment within the same cycle (Extended Fig. 10a, b, Fig. 4f). This difference in gametocyte rates was not due to delayed gametocyte maturation, as the median RFP fluorescence of gametocytes did not differ between conditions (Extended Fig. 10c). The basal gametocyte rate increased with increasing FCS or mouse serum content in the medium in a dosedependent manner up to 3-4 %, indicating that a specific molecule (or a combination) present in serum triggers same cycle sexual commitment (SSC) of early rings in *P. berghei* (Fig. 4g, Extended Fig. 10d). Notably, the gametocyte rate of normocyte-invading parasites that were cultured in rich medium remained at the basal level of 1 % (Fig. 4d), indicating that this external inducer is only active on reticulocyte-invading parasites. Reticulocytes still contain many metabolic pathways that are absent in normocytes, and it is tempting to speculate that the inducer present in serum is metabolised to its active form in reticulocytes, triggering sexual commitment in this nutrient-rich host cell only^23^. Future work will aim to identify this inducing factor. In summary, our data demonstrates that for *P. berghei* a substantial proportion of gametocytes are formed as a result of nutrient-dependent same-cycle sexual commitment triggered upon invasion of metabolically active reticulocytes.

## Conclusion

The blood stage infection in the haematopoietic niche is an emerging paradigm in malaria parasites. While extravascular maturation of gametocytes appears to be a common feature of many *Plasmodium* species^10–12^, there is much more variation in the invasion preference of asexual parasites for reticulocytes and their sequestration in extravascular erythropoietic niches^2^. Asexual replication in the erythropoietic niches can protect parasites from drug treatment, and can provide a reservoir for relapses, while the extravascular sexual development is essential for the continuation of the malaria life cycle^7^. Understanding the interactions between *Plasmodium* and the erythropoietic system should improve our ability to interfere with this extravascular parasite reservoir.

Here we present a detailed single cell analysis of host parasite interactions in erythropoietic niches using the rodent malaria model *P. berghei*. *P. berghei* exhibited a bimodal invasion pattern either into early reticulocytes or fully matured normocytes, and, in line with findings for the human reticulocyte-restricted parasite *P. vivax*^22^, the transferrin receptor CD71 served as key host receptor for reticulocyte invasion. We also found that host cell, not organ tropism determines parasite phenotypes: metabolic adaptation leads to the upregulation of the purine salvage pathway in normocytes and greatly increased sexual commitment rates in reticulocytes. Transcriptional data comparing culture-adapted *P. knowlesi* to reticulocyte-restricted *in vivo* counterparts suggests that such metabolic adaptation to the host cell is conserved across species^27^ and may represent a novel avenue for host targeted antimalarial interventions. Here we unravel complex host-parasite interactions *in vivo* on a single cell level to provide fundamental biological insights. Our study serves as a blueprint for similar investigations in the major human malaria parasites *P. falciparum* and *P. vivax*.

## Methods

### Mice

Most animal experiments were performed according to the guidelines defined by the Home Office and UK Animals (Scientific Procedures) Act 1986 and approved by the UK Home Office (project license P6CA91811) and the University of Glasgow animal welfare and ethical review body. Female TO mice weighing 25-30 g were purchased from Envigo and were aged five to eight weeks at the time point of infection. For investigating nutrient dependency of sexual commitment, animal experiments were performed according to FELASA and GV-SOLAS guidelines and approved by the responsible German authorities (Regierungspräsidium Karlsruhe). Female Swiss mice weighing 20-25 g were purchased from JANVIER and were aged five to eight weeks at the time point of infection.

### *P. berghei* infections

Most experiments were performed using a *P. berghei* line constitutively expressing mCherry under the control of the *hsp70* promoter (PbmCherry)^28^. To investigate gametocyte conversion rates in different host cells in varying nutrient conditions, we used the previously published *P. berghei* ANKA *Pb*GFP_CON_/RFP_GAM_ line that expresses GFP under the control of the constitutive ef1a promoter and RFP under the gametocyte-specific PBANKA_101270 promoter^25^. Infections were initiated by intraperitoneal infection of donor mice with ~200 μl thawed cryostocks. Depending on the experiment, recipient mice were inoculated intravenously with mixed blood stages obtained from donor mice or with synchronised schizont stages obtained as detailed below. Parasitaemia was monitored by Giemsa-stained blood smears or flow cytometry. At indicated time points post infection, mice were bled by cardiac puncture under terminal anaesthesia and culled by cervical dislocation.

### Blood and tissue collection

Blood was collected from mice under terminal anaesthesia by cardiac puncture using a heparinized syringe. For organ collection, mice were subsequently culled by cervical dislocation, spleen, femur and tibia harvested, and the organs placed into 1 ml FACS buffer (0.2 - 0.5 % BSA/PBS, sterile filtered) on ice. Bone marrow was released by removing the ends of the bones with sharp scissors and perfusing the bones with 1 ml cold FACS buffer. Spleen and bone marrow were dissociated by gently straining the tissues through a 40 μm cell strainer using the plunger of 5 ml syringe into 50 ml tubes. 50-100 μl blood were filtered through a 40 μm cell strainer into 50 ml tubes. Cell strainers were flushed with 25 ml FACS buffer and cells were pelleted by centrifugation for 10 min at 4 °C. Blood was resuspended in 20 ml, spleen in 10 ml and bone marrow in 1 ml cold FACS buffer. Cells were counted in a Neubaur counting chamber.

### Organ preparation for histology and immunofluorescence

Spleens for histological analysis were cut in half transversely and placed in 4% PFA for 5 hours, before transfer to 15% (w/v) sucrose in PBS for 2 hours and then 30% (w/v) sucrose overnight (all at 4 °C) for fixation and cryoprotection. Spleens were embedded in OCT (CellPath) using dry ice and isopentane and stored at −80°C. Sections were cut at 3-5 μm on a cryotome (ThermoFisher Scientific) using C35 carbon steel blades (Feather) on Superfrost Plus slides (ThermoFisher Scientific). Cut sections were wrapped in tin foil and stored at −80°C until staining. For Haematoxylin and Eosin (H&E) staining, sections were removed from the freezer and left wrapped in foil for 20 minutes, unwrapped and then left for a further 10 minutes at RT. Sections were fixed in ice-cold acetone/ethanol (75%/25%) for 10 minutes, air dried for 10 minutes, then placed in distilled water for 3 minutes before placement in acidified Harris haematoxylin (CellPath) for 3 minutes. Excess stain was removed by brief introduction to a running tap water bath, before two dips in 1% acetic acid in ethanol solution to differentiate, further rinsing in tap water, 30 seconds in Scott’s Tap Water substitute (Atom Scientific), one final tap water rinse, 10 dips in 70% ethanol and then 2 minutes incubation in 1% alcoholic eosin Y stain (CellPath). Slides were dehydrated gradually by incubation in 90% ethanol twice, 100% ethanol twice and finally xylene twice before mounting with a coverslip and DPX (CellPath) mountant. Slides were assessed for features from slide scans taken with a Leica Aperio Versa. 100x images were taken on an Olympus BX43F light microscope with DP22 camera (Olympus). For immunofluorescence assays, sections were removed from the freezer and left wrapped in foil for 20 minutes at RT, before rehydration in 1X PBS for 3 minutes. Sections were drawn around with a hydrophobic pen, 4% PFA added to each section and incubated in a humid chamber for 3 minutes, before washing in PBS twice for 3 minutes. Sections were permeabilised with 0.2% Triton X-100 for 10 minutes and washed again in PBS twice for 3 minutes. Sections were blocked with 2.5% normal goat serum (Vector Laboratories) supplemented with 2.5% normal mouse serum (Invitrogen) for 1 hour at RT. Sections were then incubated with 1.25 μg/ml rabbit anti-CD31 (NB100-2284 - Novus Biologicals) overnight at 4 °C. The day after, sections were rested at RT for 30 minutes before washing thrice with PBS for 3 minutes each and incubating with 1:200 goat anti-rabbit IgG-AlexaFluor647 (A21245 - Invitrogen) for 1 hour. Finally, sections were washed in PBS twice, stained with 2.5 μM DAPI (brand), washed with PBS, then treated with TrueView autofluorescence quencher (Vector Laboratories), washed PBS and mounted and coverslipped with Vectashield Vibrance (Vector Laboratories). Images were taken on a Nikon A1R Ti2 confocal microscope at 60x with immersion oil (Nikon).

### Flow cytometry analysis

Approximately 1 x 10^6^ to 1 x 10^7^ cells were transferred to a 96-well plate, pelleted for 2 min at 400 xg, resuspended in 100 μl FACS Buffer plus 5 μl TruStain fcX™ (anti-mouse CD16/32) (Biolegend Cat#101320), and incubated for 10 min on ice. Antibodies and DNA stains were added and cells were stained for 30 min on ice. Cells were pelleted (2 min at 400 xg) and washed twice with 200 μl FACS Buffer. After resuspension in 200 μl FACS Buffer, cells were analysed on a MACSQuant VYB flow cytometer or a BD FACSCelesta. Data was analysed using FlowJo software (v. 10.7.2).

### *Ex vivo* maturation of schizonts and parasite stage separation via density gradient

To obtain mature schizonts, donor mice with 1-2 % parasitaemia were bled and the blood was cultured for 16-20 hours in 25-30 ml 20 % FCS/RPMI at 5% O_2_/5% CO_2_. Schizonts were purified by density gradient centrifugation as described previously^29^. Briefly, the culture was layered on 5 ml 55 % Nycodenz/PBS, centrifuged for 20 min at 600 x g with reduced acceleration and deceleration, and schizonts were collected from the interphase. For depletion of late stages from cell suspensions, cells were separated using the same density gradient, but retaining the pellet containing ring stages and uninfected RBCSs. Either fraction was collected, washed with 10 ml 20 % FCS/RPMI, counted, and further processed as described in the individual sections.

### Parasite distribution across host organs and host cells

For the time course, mice were intravenously inoculated with 1 x 10^6^ mixed-stage iRBC. Blood and organs were harvested two, three and four days postinfection as described above. Cells were processed for flow cytometry and stained with BV421 anti-mouse CD45 (Biolegend, Cat# 103134), FITC anti-mouse CD71 (Biolegend, Cat# 113806), PE/Cy5 anti-mouse/human CD44 (Biolegend, Cat# 103010), PE/Cy7 anti-mouse Ter119 (Biolegend, Cat# 116222) at a 1:100 dilution and eBioscience™ Fixable Viability Dye eFluor™ 506 (Invitrogen, Cat# 65-0866-14) at a 1:1000 dilution.

For assessing parasite invasion, mice were intravenously inoculated with 4 x 10^6^ or 4 x 10^7^ purified schizonts. Blood and organs were harvested one hour post infection and processed for flow cytometry as described above. Cells were stained with Hoechst (abcam, ab228551) at a 1:5000 dilution and BV510 anti-mouse CD45 (Biolegend, Cat# 103137), FITC anti-mouse CD71 (Biolegend, Cat# 113806), PE/Cy5 anti-mouse/human CD44 (Biolegend, Cat# 103010), and PE/Cy7 anti-mouse Ter119 (Biolegend, Cat# 116222) at a 1:100 dilution.

### CD71 invasion assay

To obtain host cells for *in vitro* invasion, 30 to 50 μl blood were harvested from a naïve mouse, resuspended in 10 ml 0.5 % BSA/PBS, and seeded in in a 96-well plate at 5 x 10^6^ cells/well. Unspecific bindings were blocked by addition of 5 μl TruStain fcX™ (anti-mouse CD16/32) (Biolegend, Cat#101320) and incubation for 10 min on ice. Cells were stained in 50 μl total volume for at least 30 min on ice with PE/Cy7 anti-mouse Ter119 (Biolegend, Cat# 116222) (1:50 dilution) and BV510 anti-mouse CD71, clone RI7217, (Biolegend, Cat# 103145), BV510 Rat Anti-Mouse CD71, clone C2, (BD Biosciences Cat# 563112) or BV510™ Rat IgG2a, κ Isotype Ctrl Antibody (Biolegend Cat# 400553) at varying concentrations (30 μg/ml, 6 μg/ml, 1.2 μg/ml, 0.24 μg/ml, 0.048 μg/ml, 0.0096 μg/ml). Cells were pelleted (2 min at 400 x g) and resuspended in merosome-containing medium (see below).

In parallel, *Pb*mCherry schizonts were purified from an over-night culture by density gradient centrifugation as described above, counted, and resuspended in schizont medium. Merozoites were mechanically released by vigorous pipetting and sequential passage through a 5 μm filter (acrodisc) and a 1.6 μm filter (Puradisc, Whatman) as described previously^25^. Filters were flushed with schizont medium, flow-throughs pooled and diluted to 1×10^7^ ruptured schizonts/ml. 200 μl (2×10^6^ cells) per well were used to resuspend CD71-blocked host cells. After centrifugation for 10 min at 1200 x g, cells were incubated for 3 h at 37 °C, 5 % O_2_/5 % CO_2_. Cells were processed for flow cytometry as described above, staining with FITC anti-mouse CD71 (Biolegend, Cat# 113806) at a 1:100 dilution and Hoechst (abcam, ab228551) at a 1:25000 dilution.

### Separation and *Ex vivo* maturation of infected reticulocytes and normocytes

Donor mice infected with >1% *Pb*GFP_CON_/RFP_GAM_ parasites were bled and late stages including gametocytes were depleted by density gradient purification as described above. The pellet containing ring stages was washed once in 10 ml schizont medium (5 min, 500 x g), resuspended in 30 ml schizont medium and cultured for approximately 20 h on a shaker at 37 °C, 5 % O_2_/5 % CO_2_. Mature schizonts were purified by density gradient purification and 5 x 10^6^ schizonts were intravenously injected into recipient mice. One hour post infection, mice were culled and blood and spleen were collected as described above. Cell pellets were resuspended in 10 ml 0.5% BSA/PBS, leukocyte-depleted by filtering through pre-wetted Plasmodipur (Europroxima) filters and remaining late stages and gametocytes were removed by density gradient centrifugation as described above. The pellet containing freshly invaded ring stages was washed once in 10 ml MACS buffer (0.5% BSA/0.2 mM EDTA/PBS) and resuspended to 5 x 10^8^ cells/ml (blood) or 2.5 x 10^8^ cells (spleen). 400 μl cell suspension were incubated with 80 μl TruStain fcX™ (anti-mouse CD16/32) (Biolegend, Cat#101320) for 10 min on ice, followed by addition of 20 μl purified anti-mouse CD71 antibody (Biolegends, Cat# 113802) and further 30 min incubation on ice. Cells were washed with 10 ml MACS buffer (5 min at 1600 rpm) and resuspended in 800 μl MACS buffer before addition of 200 μl anti-rat IgG MicroBeads (Milteny Biotec, Cat# 130-0480501). Volumes for CD71 and microbead staining were proportionally adapted in case less than 1×10^8^ cells were obtained from the spleen. After 15 min incubation on ice and a second wash with 10 ml MACS buffer, cells were resuspended in 800 μl MACS buffer and loaded on pre-equilibrated LS columns (Milteny Biotec) placed in a magnet. The flow-through containing CD71-negative normocytes was collected and columns were washed three times with 3 ml MACS buffer, pooling all flow-throughs. CD71-positive reticulocytes were eluted by removing the columns from the magnet and flushing them with 5 ml MACS buffer using the plunger. Reticulocytes and normocytes were counted, pelleted (10 min at 1600 rpm) and resuspended in schizont medium. 2 x 10^7^ (normocytes) or 1 x 10^7^ cells (reticulocytes) were seeded in a 96-well plate and incubated for 28 h at 37 °C, 5 % O_2_/5 % CO_2_.

Aliquots of cells collected before and after MACS separation were analysed by flow as described above after staining with BV510 anti-mouse CD71 (Biolegend, #103145), PE/Dazzle-594 anti-mouse-CD45 (Biolegend, #103145), PE/Cy5 anti-mouse/human CD44 (Biolegend, #103010), PE-Cy7 anti-mouse Ter119 (Biolegend, #116222) (1:100 dilution) and Hoechst (Abcam, ab228551) (1:5000 dilution). After 28 h incubation, cells were analysed by flow cytometry after staining with PE/Dazzle-594 anti-mouse-CD45 (Biolegend, #103145), PE-Cy7 antimouse Ter119 (Biolegend, #116222) (1:100 dilution) and Hoechst (Abcam, ab228551) (1:5000 dilution). Gametocyte rates were determined by flow cytometry as percentage of RFP-positive cells among all GFP-positive parasites.

### Gametocyte conversion under varying hypoxanthine concentrations

Mature schizonts obtained from an over-night culture of *Pb*GFP_CON_/RFP_GAM_ were purified by density gradient and intravenously injected into recipient mice. Mice were bled 1 h after invasion and remaining schizonts and gametocytes removed by density gradient. The pellet was washed once in minimal medium (RPMI-1640 (#52400-025, gibco) with 0.5 % Albumax II (gibco)) and resuspended to a hematokrit of 1.25 % in minimal medium containing 250 mM hypoxanthine (c.c.pro) and 2 mM choline (Sigma Aldrich). The medium was additionally supplemented with FCS (gibco) or mouse serum obtained freshly from a naïve mouse. Parasites were cultured for 24 h at 37 °C, 5% CO_2_/5% O_2_ before analysis by flow cytometry on a BD FACSCelesta. Gametocyte rates were determined by flow cytometry as percentage of RFP-positive cells among all GFP-positive parasites.

### Cell preparation for scRNAseq

Blood, spleen and bone marrow was harvested from a naïve mouse or from an infected mouse 3 dpi with 1 x 10^6^ *Pb*mCherry parasites as described above. Dissociated spleen, bone marrow and blood cells were strained a second time through a 40 μM filter and sorted based on mCherry fluorescence using a BD FACSAria IIu or BD FACSAria III, collecting both mCherry-positive and mCherry-negative cells. Per organ, 0.5 x 10^6^ cells from each mCherry-positive and mCherry-negative fraction were pooled, pelleted for 10 min at 500 x g, 4 °C and resuspended in 100 μl 0.2 % BSA/PBS. Unspecific antibody binding was blocked by adding 10 μl TruStain fcX™ (anti-mouse CD16/32) (Biolegend, Cat#101320) to each sample and incubating for 10 min on ice. 20 μl cells were removed for flow analysis and remaining cells were incubated for 30 min on ice with 1 μg each of TotalSeq™-A0441 anti-mouse CD71 Antibody (Biolegend, Cat# 113824) and TotalSeq™-A0073 anti-mouse/human CD44 Antibody (Biolegend, Cat# 103045). Cells were washed by adding 1 ml 0.2 % BSA/PBS and pelleted for 5 min at 2800 x g, 4 °C. Pellets were resuspended in 1 ml 0.2 % BSA/PBS and washed twice more. Cells were filtered through a 40 μM strainer (Flowmi) and counted on an automatic cell counter (Countess 3, Invitrogen) immediately before loading onto the 10X Chromium machine. The 10X libraries (generated following the standard 10X protocol) were sequenced on an Illumina NextSeq 550 to ~40.000-50.000 reads per cell with reads length of 27 for read 1 and 130 for read 2. CITE-Seq libraries prepared following a previously published protocol^18^ and sequences on an Illumina NextSeq 550 with around 5000 reads per cell and CITE-Seq adapter.

### Data analysis

#### Transcriptome mapping

The raw Illumina reads were mapped with Cell Ranger (v. 3.1.0) against a combined reference of Mouse mus version 93 and Plasmodium berghei version 3 with extended 3’ UTR. CITE-Seq reads were processed with CITE-seq-Count (v. 1.4.3)

#### Data preprocessing and quality control

Quality control and data integration was performed in R (v. 3.6.1) using Seurat (v. 3.2.2)^30^. As the filtered Cell Ranger count matrices did not include RNA-low early ring stages within normocytes, the Seurat object was created from the raw count matrices including all cells with at least 100 genes and 100 UMIs. The percentage of detected mitochondrial genes in *P. berghei* (Pb) and *M. musculus* (Mm) genes and the number of UMI and genes for parasite (PbUMI and PbGenes) and host (MmUMI and MmGenes) was calculated and added as metadata. To exclude duplets, we determined upper t anvhresholds for MmGenes, MmUMI, PbGenes and PbUMI for each dataset individually by visual inspections of the plots and subsetted datasets on those. Precise thresholds per dataset are listed in Supplementary Table 1. CITE-seq information was added and barcodes missing in the CITE-seq dataset were excluded. These filtering steps retained 79.20% to 98.53% of all cells per dataset (Supplementary Table 1).

#### Host cell analysis

We analysed host cells in spleen and bone marrow by subsetting datasets on cells with more than 100 MmGenes and on features that map to *M. musculus*. Single-cell transcriptomes were normalised using scran (v. 1.14.6) including a clustering step using quickCluster() and computeSumFactors()^31^. Normalised counts were log +1 transformed. The six datasets from spleen and bone marrow (two infected, one naïve each) were integrated using the Seurat integration workflow which is based on identifying anchors across datasets to match shared cell populations^30^. Cell types were predicted by transferring labels from a bone marrow reference dataset^14^ using Seurat Transfer Labels workflow and the first 30 principal components.

For individual analysis of spleen and bone marrow cells, the combined object was subsetted per organ to cells containing at least 1000 MmUMI. Following the standard Seurat pipeline, cells were visualised using UMAP nonlinear dimensionality reduction in Seurat with the first 38 and 30 principal components for bone marrow and spleen, respectively. Clusters were identified by a shared nearest neighbour (SNN) modularity optimization based clustering algorithm using the FindClusters() function in Seurat using the default parameters and a resolution of 0.6 (bone marrow) or 0.5 (spleen). Conserved markers were identified using the Seurat built-in function with default settings and clusters were annotated according to marker gene expression, merging similar clusters.

#### Parasite cell analysis

Datasets from the six infected samples (spleen, blood and bone marrow from two mice) were subsetted on parasite-infected RBCs as defined by A) having more than 100 PbGenes and B) being predicted to be either erythroblasts (based on the label transfer in the host cell analysis) or RBCs (based on having less than 100 MmGenes). We included the malaria cell atlas data set as reference^15^. Normalisation with scran and integration of the data sets with Seurat was performed as for the host cell analysis. UMAP nonlinear dimensionality reduction was performed with the first 24 principal components and clusters were identified using a resolution of 0.51. Clusters were annotated according to correlation with the pre-annotated stages of the malaria cell atlas. One cluster with very low UMI count and ambiguous marker genes could not clearly be correlated to any parasite stage and as excluded from further analysis as outlier or duplets.

#### Differential gene expression analysis and GO enrichment

Differential gene expression analysis between host cells from infected versus uninfected mice or between *P. berghei* parasites in spleen, bone marrow and blood, or in normocytes versus reticulocytes was done using edgeR (v. 3.28.1)^32,33^. Single cells were aggregated per cluster, organ and optionally host cell by summing up counts and differential gene expression between pseudo-bulk datasets was determined using a quasi-likelihood (QL) F-test. Hits were considered significant based on false discovery rate (FDR). Due to the high amount of hits in host cell analysis, we used a more stringent cut-off for host cells (FDR < 0.01) than for *P. berghei* analysis (FDR < 0.05). Gene ontology (GO) analysis for enriched biological processes was done using the EnrichR web interphase, selecting only significantly enriched GO terms (adjusted p < 0.01) (https://maayanlab.cloud/Enrichr/)^34,35^.

### Statistics

Unless stated otherwise, experiments were repeated at least three independent times and exact sample sizes are given in the figure legends. Data was analysed and plotted using GraphPad Prism (v. 9.0.1), except for scRNA-seq data, which was analysed and plotted in R (v. 3.6.1) using Seurat built-in plotting functions or ggplot2 (v. 3.3.0). Where indicated, data was tested for significant differences using a one-way or two-way ANOVA followed by appropriate multiple comparison tests as indicated in the figure legends. Error bars indicate SEM unless stated otherwise. Significant differences between samples are indicated with asterisks as follows: *, p < 0.05; **, p < 0.01; ***, p < 0.001; ****, p < 0.0001.

## Data availability statement

The raw sequencing data and the count matrices generated and analysed during the current study will be available in the ArrayExpress repository under accession number E-MTAB-10800 (https://www.ebi.ac.uk/arrayexpress/experiments/E-MTAB-10800/). All other datasets generated and analysed during the current study are available from the corresponding author on reasonable request. Exploration of the *P. berghei* scRNA-seq data set including cluster markers and differential gene expression will be possible via a web interface upon peer-reviewed publication.

## Code availability statement

Count matrices and any code written to analyse the data is made available as GitHub repository (https://github.com/FHentzschel/MalariaHostRBCAtlas).

## Acknowledgements

We thank Craig Lapsley, Julie Galbraith and Pawel Herzyk (Glasgow Polyomics, University of Glasgow) for their guidance, library preparation and sequencing. We thank Chris J. Janse (Leiden University Medical Center) for providing the PbmCherry line and Andrew P. Waters (University of Glasgow) for providing *Pb*GFP_CON_/RFP_GAM_. We thank Manoj Duraisingh (Harvard T.H. Chan School of Public Health) for help with setting up the CD71 invasion assay and Katarzyna Modrzynska (University of Glasgow) for experimental advice and assistance. We thank Meng Du for exploration of pseudo time methods for the dataset. F.H. was funded by the Deutsche Forschungsgemeinschaft (DFG, German Research Foundation), project number 404044656. M.M. is funded by a Wolfson Merit Royal Society Award. This work was funded through Wellcome Trust Investigator award 110166, Wellcome Trust Center award 104111, ERC Consolidator Award BoneMalar to M.M and through Wellcome Trust (no. 204820/Z/16/Z), to T.D.O.

## Author contributions

F.H., M.M. and T.D.O. conceived the study and designed the experiments. M.M., C.A.M. and T.D.O. supervised the work. F.H. performed most experiments and data analysis, with assistance by M.P.G. M.P.G. performed histology experiments. C.A. performed histopathological analysis. F.H., D.B. and T.D.O. analysed the scRNA-seq data. F.H. and M.M. wrote the manuscript with input and comments from all authors.

## Additional information

Supplementary Information is available for this paper.

## References

1. World Health Organization. WHO | The World malaria report 2020. https://www.who.int/publications/i/item/9789240015791 (2020).

2. Venugopal, K., Hentzschel, F., Valkiūnas, G. & Marti, M. Plasmodium asexual growth and sexual development in the haematopoietic niche of the host. Nature Reviews Microbiology 18, 177–189 (2020).

3. Kho, S. et al. Evaluation of splenic accumulation and colocalization of immature reticulocytes and Plasmodium vivax in asymptomatic malaria: A prospective human splenectomy study. PLOS Medicine 18, e1003632 (2021).

4. Kho, S. et al. Hidden Biomass of Intact Malaria Parasites in the Human Spleen. New England Journal of Medicine 384, 2067–2069 (2021).

5. Malleret, B. et al. Significant Biochemical, Biophysical and Metabolic Diversity in Circulating Human Cord Blood Reticulocytes. PLoS ONE 8, e76062 (2013).

6. Chen, K. et al. Resolving the distinct stages in erythroid differentiation based on dynamic changes in membrane protein expression during erythropoiesis. Proceedings of the National Academy of Sciences of the United States of America 106, 17413–8 (2009).

7. Lee, R. S., Waters, A. P. & Brewer, J. M. A cryptic cycle in haematopoietic niches promotes initiation of malaria transmission and evasion of chemotherapy. Nature Communications 9, 1689 (2018).

8. Neveu, G. et al. Plasmodium falciparum sexual parasites develop in human erythroblasts and affect erythropoiesis. Blood 136, 1381–1393 (2020).

9. Peatey, C. L. et al. Enhanced gametocyte formation in erythrocyte progenitor cells: A site-specific adaptation by plasmodium falciparum. Journal of Infectious Diseases 208, 1170–1174 (2013).

10. Niz, M. de et al. Plasmodium gametocytes display homing and vascular transmigration in the host bone marrow. Science advances 4, 1–16 (2018).

11. Obaldia, N. et al. Bone marrow is a major parasite reservoir in plasmodium vivax infection. mBio 9, e00625–18 (2018).

12. Joice, R. et al. Plasmodium falciparum transmission stages accumulate in the human bone marrow. Science translational medicine 6, 244re5 (2014).

13. Aguilar, R. et al. Molecular evidence for the localization of plasmodium falciparum immature gametocytes in bone marrow. Blood 123, 959–966 (2014).

14. Baccin, C. et al. Combined single-cell and spatial transcriptomics reveal the molecular, cellular and spatial bone marrow niche organization. Nature Cell Biology 22, 38–48 (2020).

15. Howick, V. M. et al. The malaria cell atlas: Single parasite transcriptomes across the complete Plasmodium life cycle. Science 365, eaaw2619 (2019).

16. Hliscs, M., Nahar, C., Frischknecht, F. & Matuschewski, K. Expression Profiling of Plasmodium berghei HSP70 Genes for Generation of Bright Red Fluorescent Parasites. PLoS ONE 8, e72771 (2013).

17. Safeukui, I. et al. Retention of Plasmodium falciparum ring-infected erythrocytes in the slow, open microcirculation of the human spleen. Blood 112, (2008).

18. Stoeckius, M. et al. Simultaneous epitope and transcriptome measurement in single cells. Nature Methods 14, 865–868 (2017).

19. Zheng, G. X. Y. et al. Massively parallel digital transcriptional profiling of single cells. Nature Communications 8, 1–12 (2017).

20. Schaum, N. et al. Single-cell transcriptomics of 20 mouse organs creates a Tabula Muris. Nature 562, 367–372 (2018).

21. Franke-Fayard, B., Fonager, J., Braks, A., Khan, S. M. & Janse, C. J. Sequestration and Tissue Accumulation of Human Malaria Parasites: Can We Learn Anything from Rodent Models of Malaria? PLoS Pathogens 6, e1001032 (2010).

22. Gruszczyk, J. et al. Transferrin receptor 1 is a reticulocytespecific receptor for Plasmodium vivax. Science 359, 48–55 (2018).

23. Srivastava, A. et al. Host Reticulocytes Provide Metabolic Reservoirs That Can Be Exploited by Malaria Parasites. PLOS Pathogens 11, e1004882 (2015).

24. van Dijk, M. R. et al. Three Members of the 6-cys Protein Family of Plasmodium Play a Role in Gamete Fertility. PLoS Pathogens 6, e1000853 (2010).

25. Lee, R. S., Waters, A. P. & Brewer, J. M. A cryptic cycle in haematopoietic niches promotes initiation of malaria transmission and evasion of chemotherapy. Nature Communications 9, 1689 (2018).

26. Niikura, M. et al. G-strand binding protein 2 is involved in asexual and sexual development of Plasmodium berghei. Parasitology International 76, 102059 (2020).

27. Lapp, S. A. et al. Plasmodium knowlesi gene expression differs in ex vivo compared to in vitro blood-stage cultures. Malaria Journal 14, 110 (2015).

28. Burda, P.-C. et al. A Plasmodium Phospholipase Is Involved in Disruption of the Liver Stage Parasitophorous Vacuole Membrane. PLOS Pathogens 11, e1004760 (2015).

29. Janse, C. J., Ramesar, J. & Waters, A. P. High-efficiency transfection and drug selection of genetically transformed blood stages of the rodent malaria parasite Plasmodium berghei. Nature Protocols 1, 346–356 (2006).

30. Stuart, T. et al. Comprehensive Integration of Single-Cell Data. Cell 177, 1888–1902.e21 (2019).

31. Lun, A. T. L., McCarthy, D. J. & Marioni, J. C. A step-by-step workflow for low-level analysis of single-cell RNA-seq data with Bioconductor. F1000Research 5, 2122 (2016).

32. Robinson, M. D., McCarthy, D. J. & Smyth, G. K. edgeR: A Bioconductor package for differential expression analysis of digital gene expression data. Bioinformatics 26, 139–140 (2009).

33. McCarthy, D. J., Chen, Y. & Smyth, G. K. Differential expression analysis of multifactor RNA-Seq experiments with respect to biological variation. Nucleic Acids Research 40, 4288–4297 (2012).

34. Kuleshov, M. v. et al. Enrichr: a comprehensive gene set enrichment analysis web server 2016 update. Nucleic acids research 44, W90–W97 (2016).

35. Chen, E. Y. et al. Enrichr: Interactive and collaborative HTML5 gene list enrichment analysis tool. BMC Bioinformatics 14, 1–14 (2013).

