## Extended Figures for "Host cell maturation modulates parasite invasion and sexual differentiation in *Plasmodium*"

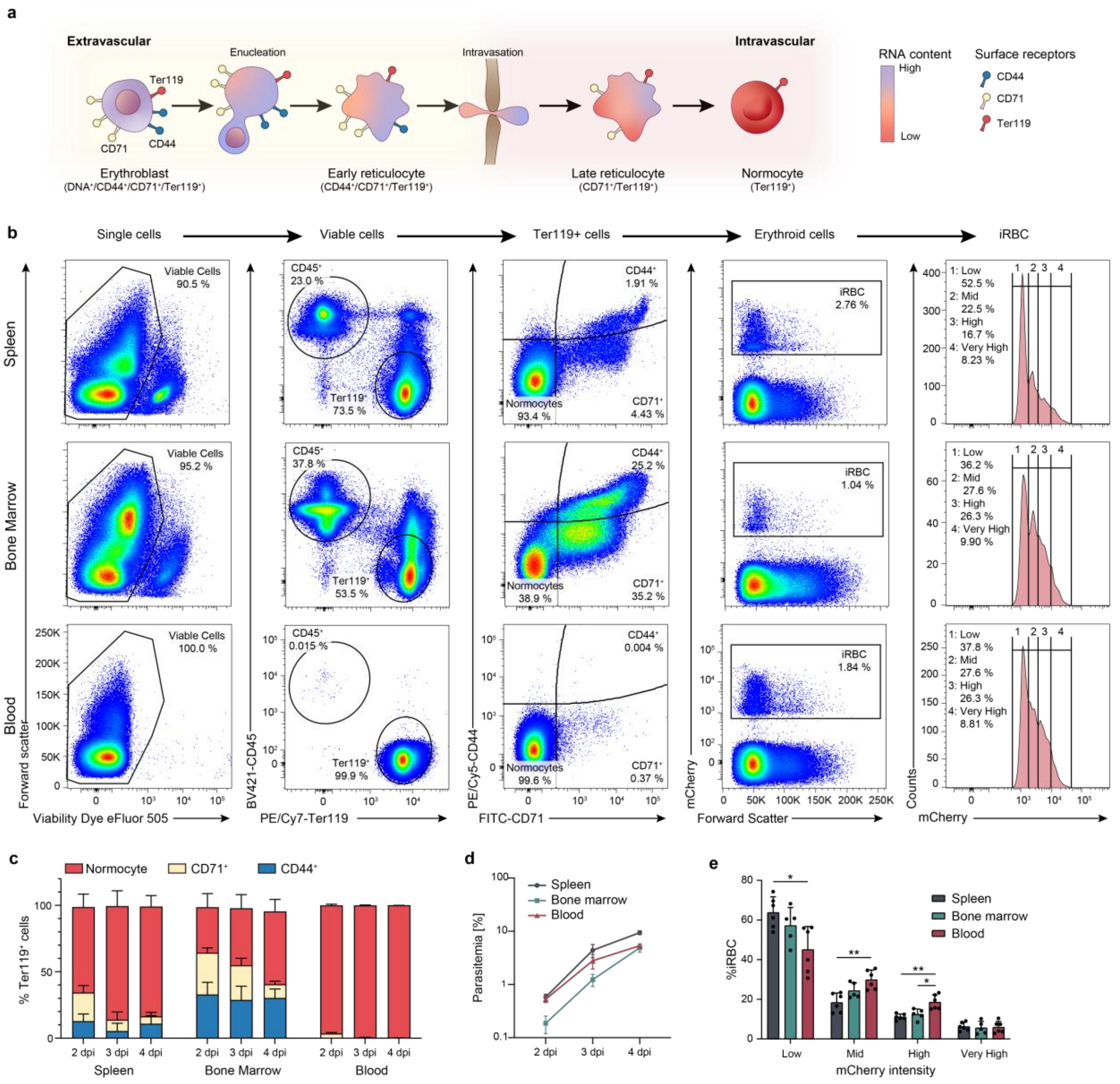


Extended Figure 1: **Time course analysis of *P. berghei* infection in spleen, bone marrow and blood**. **a**, Schematic depiction of erythropoiesis, including surface receptors indicative of erythroblasts, extravascular and intravascular reticulocytes and normocytes. **b**, Gating strategy and exemplary flow plots for time course across organs. **c**, Distribution of host RBCs across organs and time points (n = 6, except for 4 dpi, where n =3). **d**, Parasitemia over time in different organs (n = 6, except for 4 dpi, where n =3). **e**, Percentage of parasites per mCherry gate per organ as proxy for parasite stages 2 dpi. (n = 6. 2-way ANOVA, Tukey's post test, *, $p < 0.05$, **, $p < 0.01$). **c**, **d**, **e**, mean +/- SEM.

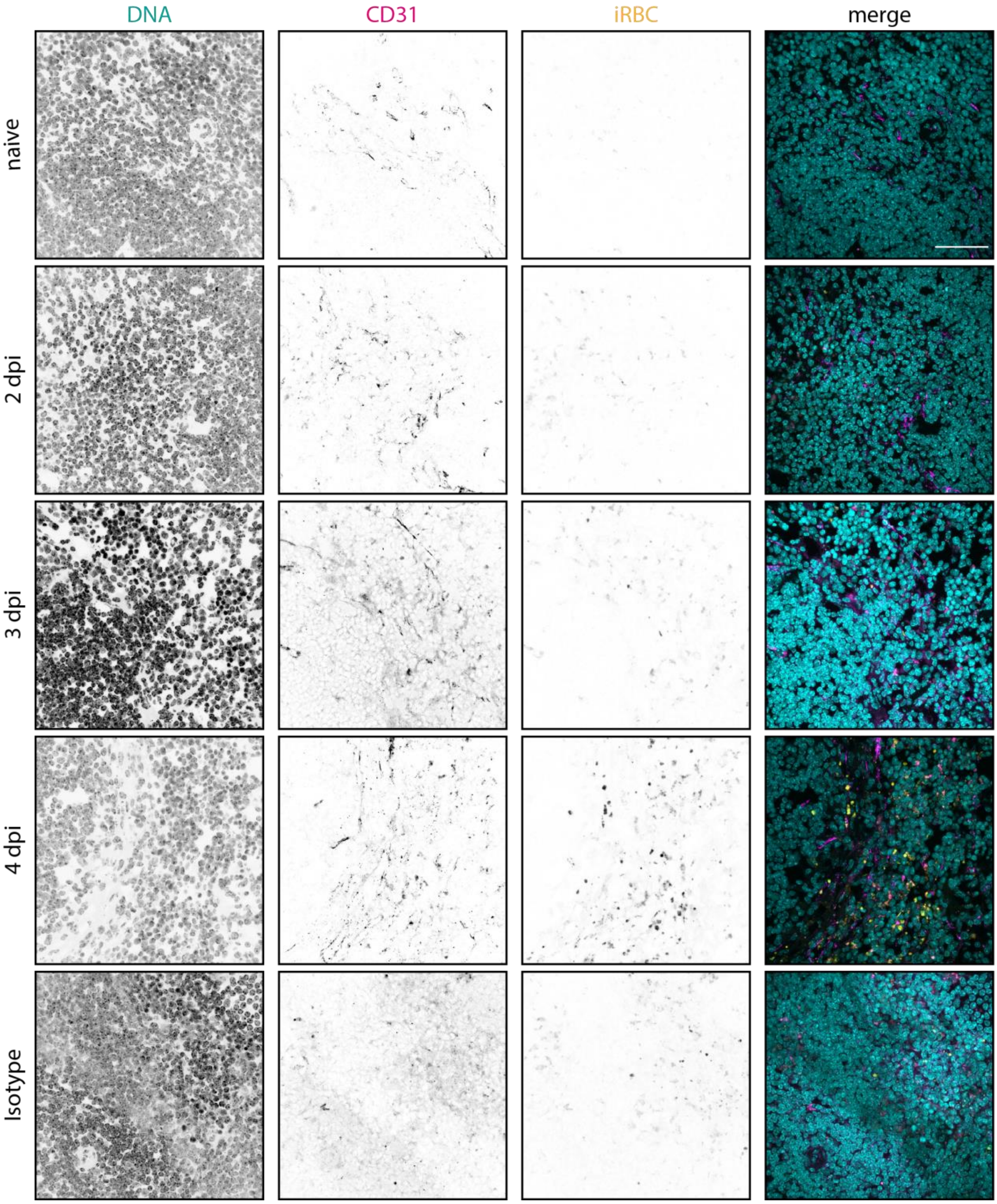


Extended Figure 2: ***P. berghei* localisation in spleen.** Representative immunofluorescence images of *P. berghei* (pseudocolored in yellow) in spleen sections stained for the endothelial cell marker CD31 (magenta). Bottom row: Isotype control, no CD31 primary antibody. Scale bar 50 µm.

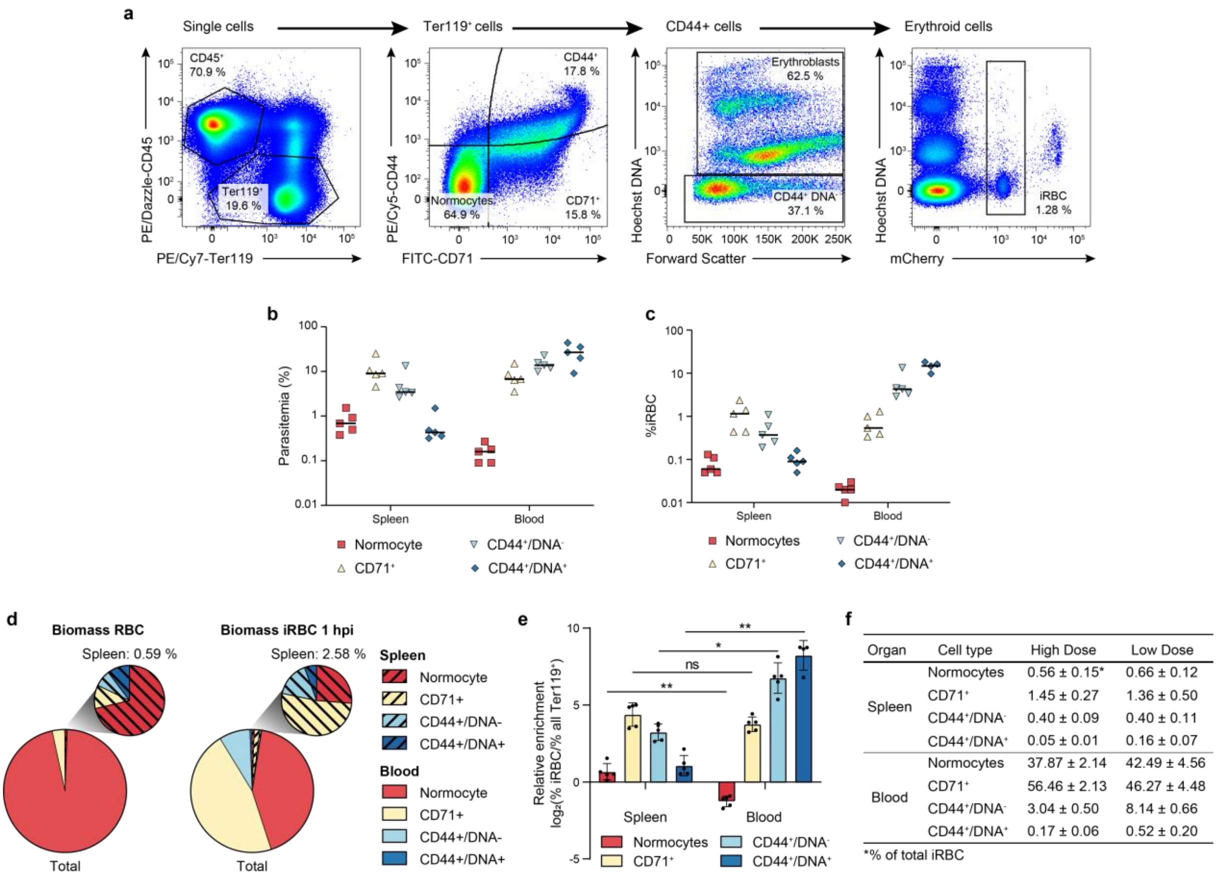


| Organ | Cell type | High Dose | Low Dose |
|---|---|---|---|
| Spleen | Normocytes | 0.56 ± 0.15* | 0.66 ± 0.12 |
| | $CD71^+$ | 1.45 ± 0.27 | 1.36 ± 0.50 |
| | $CD44^+/DNA^-$ | 0.40 ± 0.09 | 0.40 ± 0.11 |
| | $CD44^+/DNA^+$ | 0.05 ± 0.01 | 0.16 ± 0.07 |
| Blood | Normocytes | 37.87 ± 2.14 | 42.49 ± 4.56 |
| | $CD71^+$ | 56.46 ± 2.13 | 46.27 ± 4.48 |
| | $CD44^+/DNA^-$ | 3.04 ± 0.50 | 8.14 ± 0.66 |
| | $CD44^+/DNA^+$ | 0.17 ± 0.06 | 0.52 ± 0.20 |

*% of total iRBC

Extended Figure 3: ***P. berghei*** **invasion preference across organs**. **a**, Gating strategy to assess ring-stage parasite distribution 1 hpi. **b**, **c**, Parasitemia in splenic and blood RBC cell types one hour post i.v. infection with (**b**) $4*10^7$ or (**c**) $4*10^6$ iRBC. (n = 5, line at mean). **d**, Biomass of RBC cell types (left) and iRBC cell types 1 hpi with $4*10^6$ iRBC (right) in total RBCs (large pie chart) and spleen RBCs (small pie chart) (n = 5). **e**, Relative enrichment of host cell types in iRBC compared to all RBC 1 hpi with $4*10^6$ iRBC. Values > 0 indicate preferential invasion of this host cell type, values < 0 indicate decreased invasion. (n = 5. 2-way ANOVA, Sidáks post test. mean +/- SEM. ns, not significant, **, $p < 0.01$). **f**, Summary table depicting ring stage biomass ± SEM across organs and host cells 1 hpi with low ($4* 10^6$ iRBC) or high ($4*10^7$ iRBC) dose. (n = 5).

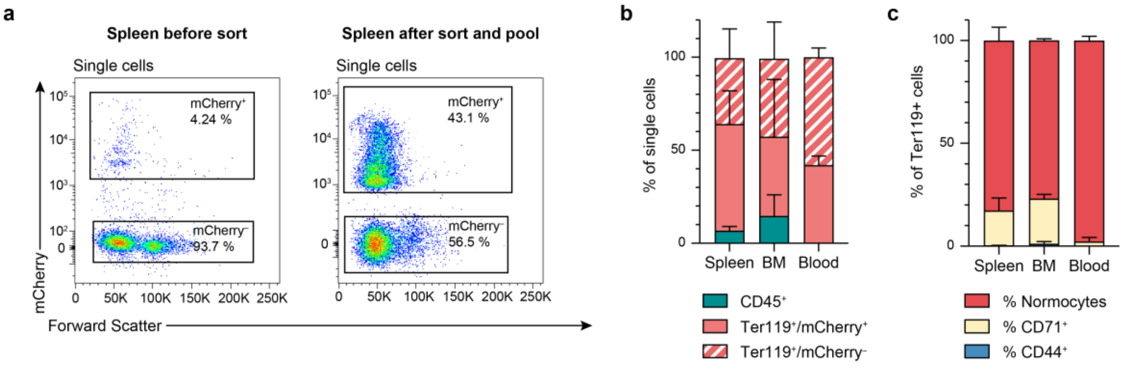


Extended Figure 4: **Flow sorting of iRBCs and host cells prior to scRNA-seq**. **a**, Exemplary flow plot showing spleen sample and sorting gates before (left panel) and after sorting and 1:1 pooling of mCherry+ iRBC and mCherry- host cells (right panel). **B**, Cell type distribution in scRNA-seq samples. **c**, RBC distribution in Ter119+ cells in scRNAseq samples. **b**, **c**: n = 2, Mean +/- SEM.

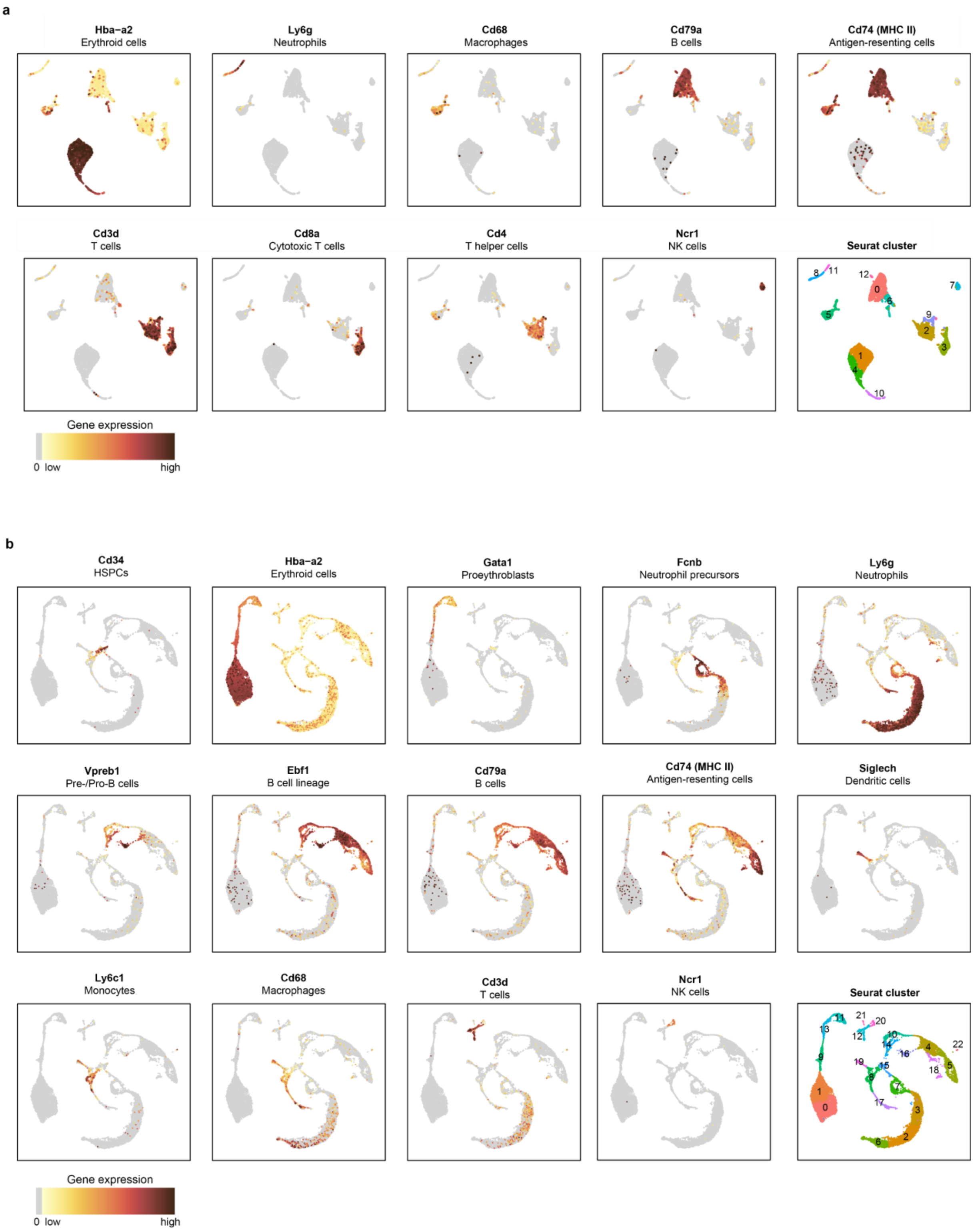


Extended Figure 5: **Marker gene expression in host cells**. **a**, **b**, UMAP of (**a**) spleen and (**b**) bone marrow colored by expression strength of canonical marker genes or according to the original clusters as identified by the unsupervised clustering approach (last panel).

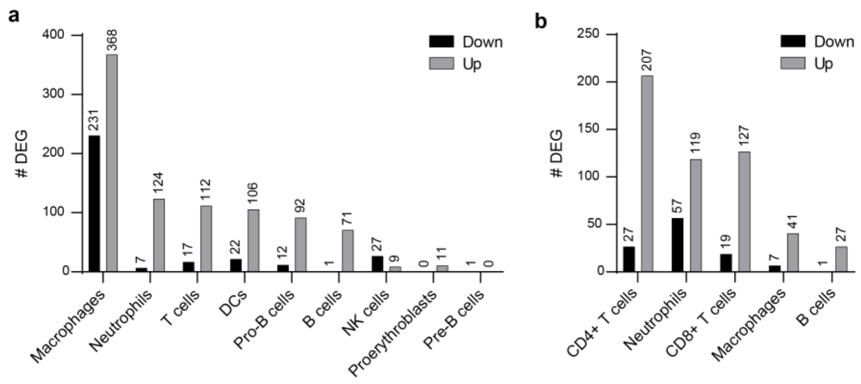


Extended Figure 6: **Host cell response to infection**. **a**, **b**, Total number of significantly differentially expressed genes (DEG) upon infection in each cell cluster of (**a**) bone marrow and (**b**) spleen.

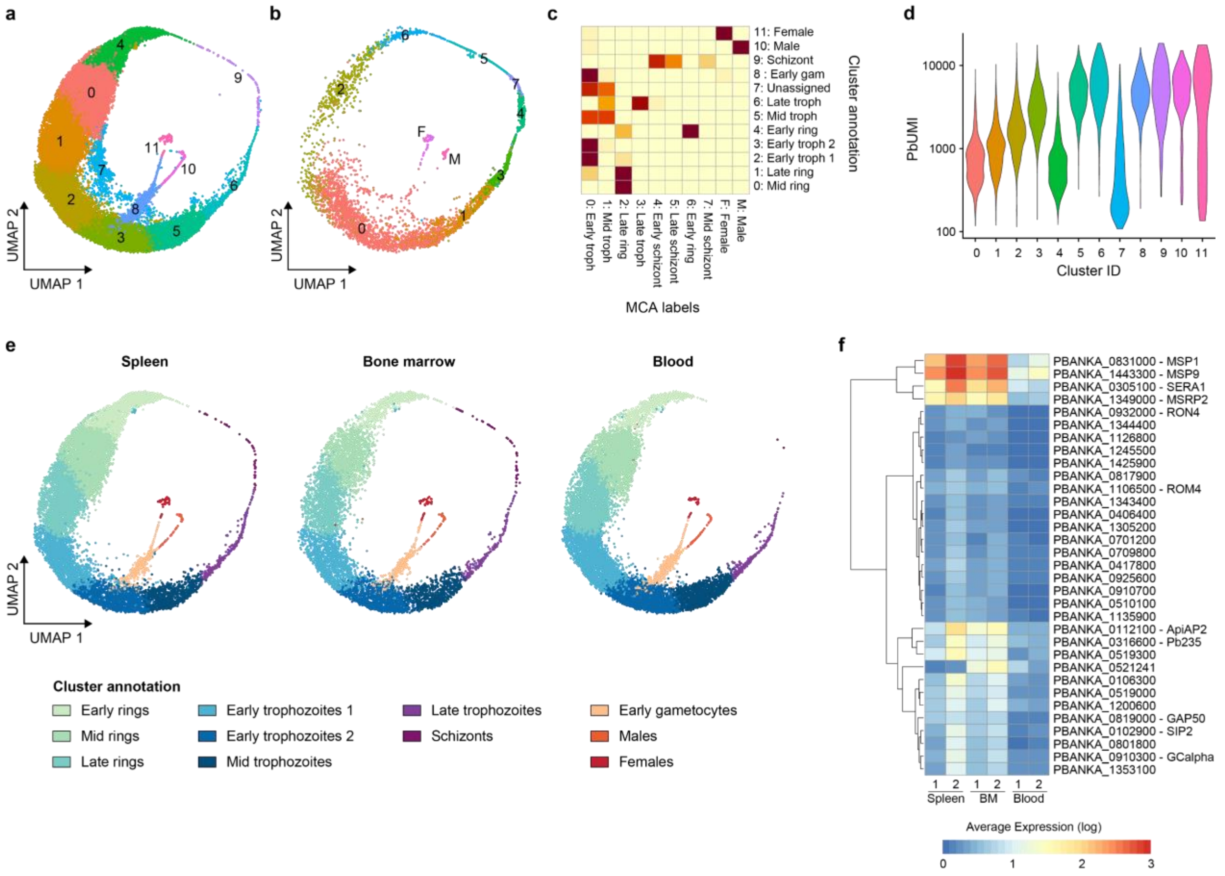


Extended Figure 7: **scRNA-seq analysis of *P. berghei* cells. a**, UMAP of P. berghei single cell transcriptomes generated in this study, colored by cluster. **b**, UMAP of malaria cell atlas. Cells are colored by their annotation in the malaria cell atlas[1]. The data set was merged with the here generated *P. berghei* data and analysed in parallel. **c**, Heatmap depicting correlation between MCA annotation and clusters. Clusters were annotated based on the MCA labels they correlated with, and the annotation is indicated next to the cluster number. **d**, *Pb*UMI per cluster ID. Cluster 7 shows a very low UMI per cell and was excluded from further analysis. **e,** UMAP of *P. berghei* cells separated according to host organ and colored according to stage. **f**, Average expression of genes in late trophozoite clusters per sample. Depicted are the most significant hits that are differentially expressed between organs in the late trophozoite cluster. Gene names of annotated genes are listed next to the Gene ID.

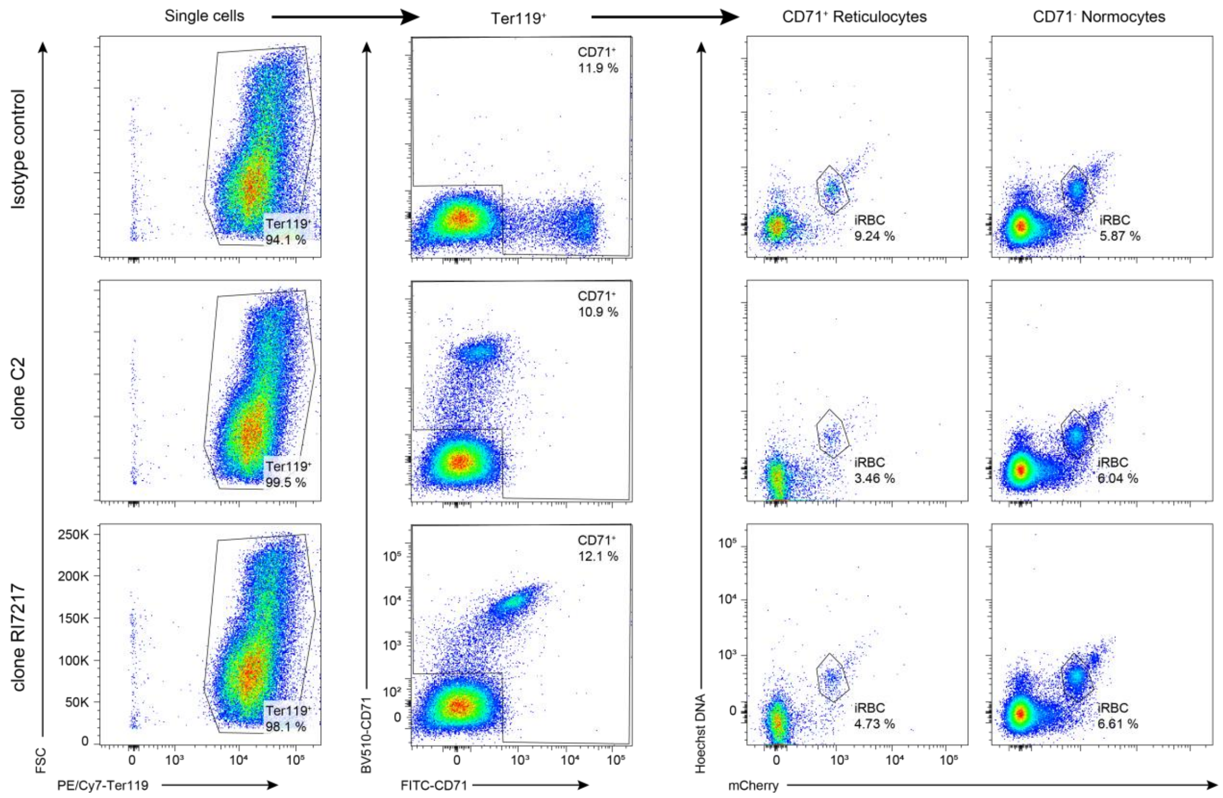


Extended Figure 8: **Gating strategy of CD71-dependent invasion assay**. RBCs were pre-incubated with isotype control or one of two clones of anti-CD71 antibodies (clone C2 or RI7217, both BV510-coupled) before invasion of *P. berghei merozoites*. After invasion, RBCs were co-stained with a second CD71 antibody coupled to FITC and invasion in $CD71^+$ reticulocytes and $CD71^-$ normocytes was determined by flow cytometry.

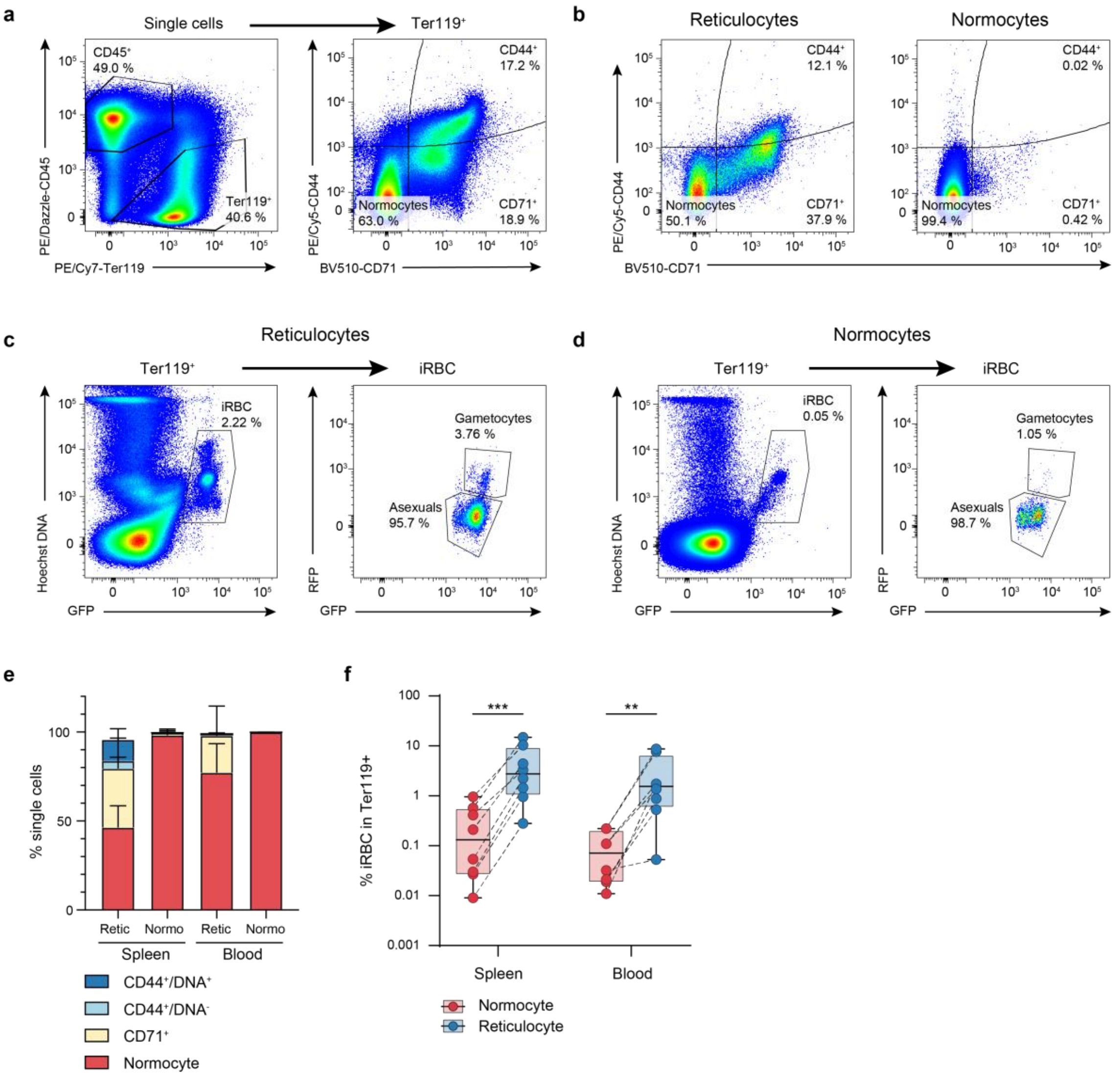


Extended Figure 9: ***Ex vivo* gametocyte conversion in reticulocytes versus normocytes**. **a**, **b**, Exemplary flow plot and gating strategy of cells (**a**) before and (**b**) after magnetic cell sorting, yielding a CD71$^+$ enriched reticulocyte fraction and a CD71$^-$ depleted normocyte fraction. **c**, **d,** Gating strategy for asexual and sexual parasitemia 28 hpi in (**c**) reticulocyte and (**d**) normocyte fractions. **e**, Host cell distribution in different fractions directly after magnetic cell sorting. (n= 8). **f**, Parasitemia after 28 h *ex vivo* culture of normocyte- or reticulocyte-enriched iRBC isolated from spleen or blood. Samples from one organ were matched across host cell. (n = 8. Two-way ANOVA, Tukey's post test).

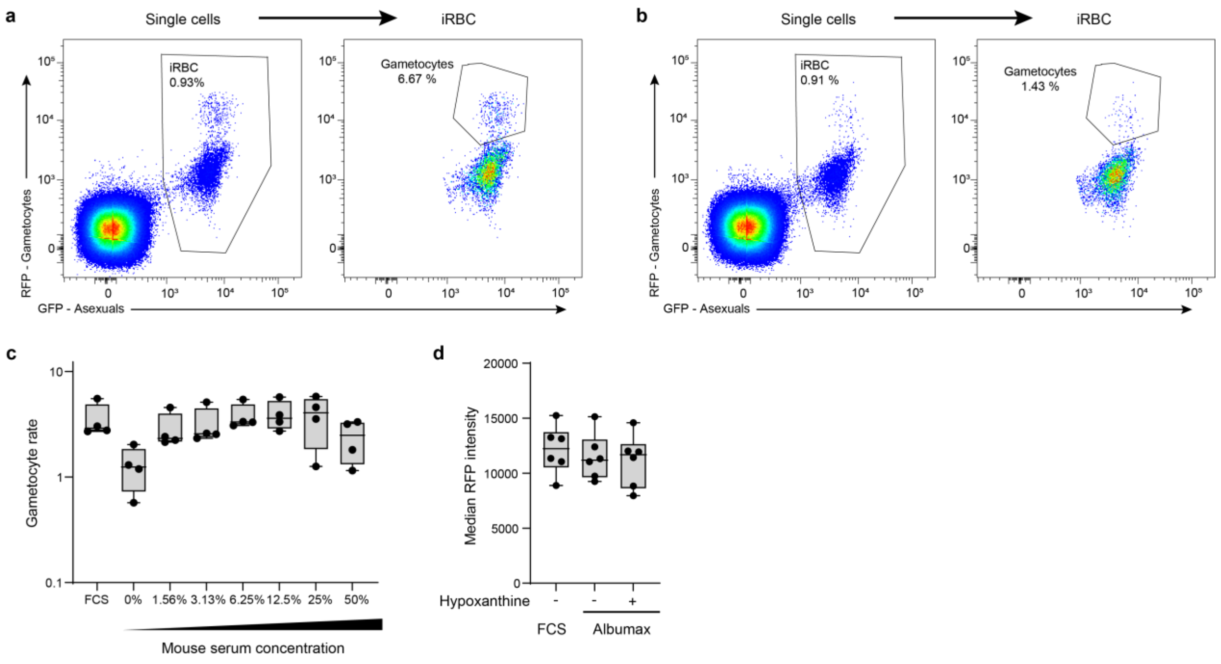


Extended Figure 10**: Nutrient-dependent same-cycle sexual conversion rate**. **a**, **b,** Gating strategy to determine sexual conversion in different media. **A,** Culture in 20% FCS. **B**, Culture in 0.5% Albumax/choline. **c,** Gametocyte rate after *ex vivo* culture in minimal medium supplemented with varying concentrations of mouse serum. (n = 4. One-way ANOVA, Dunnett's post test. *: p value < 0.05). **d**, Median RFP intensity of gametocytes in FCS/RPMI or Albumax/RPMI.
